# A programmable pAgo nuclease with RNA target preference from the psychrotolerant bacteria *Mucilaginibacter paludis*

**DOI:** 10.1101/2021.06.08.447469

**Authors:** Wenqiang Li, Yang Liu, Fei Wang, Lixin Ma

## Abstract

Argonaute (Ago) proteins are programmable nuclease found in both eukaryotes and prokaryotes. Prokaryotic Argonaute proteins (pAgos) share a high degree of structural homology with eukaryotic Argonaute proteins (eAgos) and eAgos are considered to evolve from pAgos. However, the majority of studied pAgos prefer to cleave DNA targets, and eAgos exclusively cleave RNA targets. Here, we characterize a novel pAgo, *Mbp*Ago, from psychrotolerant bacteria *Mucilaginibacter paludis* that can be programmed with DNA guides and prefers to cleave RNA targets rather than DNA targets. *Mbp*Ago can be active at a wide range of temperatures (4-65°C). In comparison with previously studied pAgos, *Mbp*Ago is able to utilize 16-nt long 5’phosphorylated and 5’hydroxylated DNA guides for efficient and precise cleavage and displays no obvious preference for the 5’end nucleotide of a guide. Furthermore, the cleavage efficiency can be regulated by mismatches in the central and 3’supplementary regions of the guide. *Mbp*Ago can efficiently cleave highly-structured RNA targets using both 5’phosphorylated and 5’hydroxylated DNA guides in the presence of Mg^2+^ or Mn^2+^. In conclusion, we have demonstrated that *Mbp*Ago is a unique programmable nuclease that has a strong preference for RNA targets, with great potential applications in the field of nucleic acid biotechnology.

## INTRODUCTION

Argonaute proteins (Agos) are present in all three domains of life, and they use small DNA or RNA guides for recognition and sometimes cleavage of complementary nucleic acid targets (1). As the key components of RNA interference (RNAi) pathways, eukaryotic Argonautes (eAgos) participate in post-transcriptional gene regulation and antivirus defence (2). Similarly to eAgos, prokaryotic Argonautes (pAgos) have been shown to protect cells from mobile genetic elements, including plasmids and phages (3–6). In addition, a recent study showed that pAgos may function in the DNA replication (7), which suggests that the cellular functions of pAgos are diverse and remains to be further explored.

Genomic studies showed that pAgos are more diverse than eAgos (8), and pAgos can be classified into long pAgos, short pAgos and PIWI-RE (9). Despite these divergent cellular functions and diversity, pAgos and eAgos adopt a highly conserved six domains architecture, including the N-terminal (N), linker 1 (L1), middle (MID), linker 2 (L2), PIWI-Argonaute-Zwille (PAZ), and P element-induced wimpy testis (PIWI) domains (5). Molecular structures of Agos showed that the 5’ end and the 3’ end of nucleic acid guides are anchored in the MID and PAZ domains, respectively (10,11). Agos with endonuclease activity have a catalytic tetrad DEDX (X is D, H, N, or K) in the PIWI domain, which is essential for binding divalent metal ions and responsible for catalysis (5,12,13). After binding with small nucleic acid guides, active Agos can precisely cleave the complementary targets between the 10’ and 11’ nucleotides of the guide nucleic acid (14).

While eAgos exclusively cleave RNA targets, previously characterized pAgos prefer to cleave DNA targets. Initially, pAgos were characterized from thermophilic prokaryotes, including *Tt*Ago (*Thermus thermophilus*) (3), *Pf*Ago (*Pyrococcus furiosus*) (4), *Mp*Ago (*Marinitoga piezophile*) (13), and *Mj*Ago (*Methanocaldococcus jannaschii*) (15,16), which can be programmed with DNA guides or RNA guides to cleave DNA targets or RNA targets effectively in elevated temperatures but not in moderate temperatures. Recently, active pAgos from mesophilic prokaryotes including *Cb*Ago (*Clostridium butyricum*) (17,18), *Lr*Ago (*Limnothrix rosea*) (17), *Se*Ago (*Synechococcus elongatus*) (19), *Km*Ago (*Kurthia massiliensis*) (20,21), *Cp*Ago (*Clostridium perfringens*) and *Ib*Ago (*Intestinibacter bartlettii*) (22) were characterized to search for a pAgo that can cleave double-stranded DNA (dsDNA) at moderate temperatures and be used for genome editing. Besides, some pAgos perform guide-independent target DNA processing activity, including *Tt*Ago, *Mj*Ago, *Cb*Ago, *Lr*Ago, and *Se*Ago (15,17,19,23). Among them, *Km*Ago can utilize both DNA and RNA guides to cleave almost all types of nucleic acids at moderate temperatures, including single-stranded DNA (ssDNA), dsDNA, unstructured RNA and highly-structured RNA, but can only use 5’phosphorylated DNA (5’P-DNA) guide to cleave RNA with high efficiency (20).

The programmable binding and cleavage activities of pAgos are similar to that of the CRISPR-Cas system, which means pAgos can potentially be used in genome editing applications (9), molecular cloning (24), and nucleic acid detection (25–28). Though eAgos prefer to use small RNA guides to cleave RNA targets, there are only a few pAgos that have been reported to recognize RNA targets, and no pAgos with the ability to preferentially cleave RNA targets with high accuracies at mesophilic temperatures have been described to date. The search for pAgos capable of cleave RNA targets with high specificity and high efficiency at 37°C, which could have widespread applications in RNA fields, is therefore of great importance. In this work, we characterize a novel pAgo, *Mbp*Ago from *Mucilaginibacter paludis*, which is distantly related to other characterized pAgos, and contains the canonical catalytic tetrad in the PIWI domain (residues D566, E601, D635, and D768). We show that different from other pAgos, *Mbp*Ago binds DNA guides to cleave RNA with high efficiency but DNA with very low efficiency at physiological temperatures. Furthermore, we demonstrated that *Mbp*Ago can utilize both 5’ phosphorylated and 5’ hydroxylated DNA guides to cleave unstructured RNA and highly-structured RNA, which suggests that it can expand the toolkit for RNA-manipulation and detection.

## MATERIAL AND METHODS

### Protein expression and purification

The *Mbp*Ago gene (WP_008504757.1; *Mucilaginibacter paludis)* and *Mbp*Ago double mutant (*Mpb*Ago_DM) (D566A, D635A) gene were synthesized by Wuhan Genecreate Biotechnology Co., Ltd and cloned into pET28a expression vectors. The *Mbp*Ago and *Mbp*Ago_DM proteins were expressed in *E. coli* Rosetta (DE3) (Novagen). Cultures were grown at 37 °C in Luria-Bertani (LB) medium containing 50 μg/ml kanamycin and until OD600 reached 0.8. *Mbp*Ago expression was induced by the addition of isopropyl-β-D-1-thiogalactopyranoside (IPTG) to a final concentration of 0.5 mM. Cells were incubated at 18°C for 16 h with continuous shaking for expression. Centrifugally collected cells were stored at a −80°C refrigerator for further protein purification.

The cell pellet was resuspended in Buffer A (20 mM Tris-HCl pH 7.4, 500 mM NaCl, 20 mM imidazole) supplemented with 1 mM of phenylmethylsulfonyl fluoride (PMSF) and disrupted by sonication (SCIENTZ-IID. 400 W, 2 s on/4 s off for 15 min). The lysate was clarified by centrifugation and the supernatant was loaded onto Ni-NTA agarose resin for 50 min with rotation. The beads were washed with Buffer A, then with the same buffer containing 50 mM imidazole and eluted with Buffer A containing 150 mM imidazole. Fractions containing *Mbp*Ago were concentrated by ultrafiltration using an Amicon 50K filter unit (Millipore) and purified on a Superdex 200 16/600 column (GE Healthcare) equilibrated with Buffer B (20 mM Tris-HCl pH 7.4, 500 mM NaCl). Fractions containing *Mbp*Ago were concentrated Amicon 50K filter unit (Merck Millipore), placed in Buffer B (20 mM Tris-HCl pH 7.4, 500 mM NaCl), aliquoted and flash-frozen in liquid nitrogen.

### Single-stranded nucleic acid cleavage assays

The cleavage assays were performed using the synthetic guide and target (see Supplementary Table S1 for oligonucleotide sequences). 5’-FAM-labeled targets and 5’-P-labeled guides were synthesized for some experiments. The cleavage reactions were performed in PCR tubes at 37 °C in a buffer containing 10 mM HEPES-NaOH pH 7.5, 100 mM NaCl, 5 mM MnCl2, and 5% glycerol. To analyze the effect of various divalent cations, 5 mM Mg^2+^, Ni^2+^, Co^2+^, Cu^2+^, Fe^2+^, Ca^2+^, or Zn^2+^ were added instead of Mn^2+^. 800 nM of *Mbp*Ago was mixed with 400 nM DNA guide or RNA guide and incubated for 10 min at 37 °C for guide loading. Target nucleic acids were then added to the final concentration of 200 nM. For analysis of temperature dependence of RNA cleavage, *Mbp*Ago was loaded with DNA guide for 10 min at 37°C, the samples were transferred to indicated temperatures in a PCR thermocycler (T100, Bio-Rad), RNA target was added and the samples were incubated for 15 min. All reactions were carried out at 37°C if not indicated. The reactions were stopped after indicated time intervals by mixing the samples with equal volumes 2× RNA loading dye (95% Formamide, 18 mM EDTA, and 0.025% SDS, and 0.025% Bromophenol Blue) and heated for 5 min at 95°C. The cleavage products were resolved by 20% denaturing PAGE, stained with SYBR Gold (Invitrogen), visualized with Gel Doc™ XR+ (Bio-Rad), and analyzed by the ImageJ and Prism 8 (GraphPad) software. For reactions containing FAM labels, the gels were first visualized with Gel Doc™ XR+ (BioRad) and then stained with SYBR Gold.

### Highly-structured RNA cleavage assays

The SARS-CoV-2 RdRp partial RNA (see Supplementary Table S2) were *in vitro* transcribed using T7 RNA polymerase (Thermo Fisher Scientific) and synthetic DNA templates carrying a T7 promoter sequence. The transcripts were treated with DNase I, gel-purified, and ethanol precipitated. DNA guides used for cleavage (Supplementary Table S3) were 5’-phosphorylated using T4 PNK (New England Biolabs) except when cleavage with 5’-OH guides. 1 μM of *Mbp*Ago was mixed with 1 μM guide DNA and incubated for 10 min at 37 °C for guide loading. Target RNA was then added to the final concentration of 250 nM and incubated for 30 min at 37°C for cleavage. Reactions were stopped with 2× RNA loading dye and heated for 5 min at 95°C. The cleavage products were resolved by 8% denaturing PAGE and stained with SYBR Gold.

### Electrophoretic mobility shift assay (EMSA)

To examine the loading of guide onto *Mbp*Ago, *Mbp*Ago and 3’end FAM-labeled guide were incubated in 10 μl of reaction buffer (10 mM HEPES-NaOH pH 7.5, 100 mM NaCl, 5mM Mn^2+^) for 30 min at 37°C. The concentration of the guide was fixed as 100 nM, whereas the concentration of *Mbp*Ago varied. Then the samples were mixed with 1 μl 10× loading buffer (250 mM Tris-HCl (pH 7.5), 40% glycerol) and resolved by 10% native PAGE with 0.5× Tris-Borate-EDTA (TBE) buffer. Nucleic acids were visualized using Gel Doc™ XR+. To analyze the loading of the guide:target duplex, reactions contained *Mbp*Ago_DM, 100 nM 3’end FAM-labeled guide, 100 nM target, were combined for 10 μl and incubated for 30 min at 37°C. The concentration of *Mbp*Ago_DM varied. Then the samples were mixed with 1 μl 10× loading buffer and resolved by 10% native PAGE with 0.5× TBE. Nucleic acids were visualized using Gel Doc™ XR+. To determine the apparent dissociation constants (Kd) for guide binding and guide:target duplex binding, the obtained gel images were analyzed with the NIH program ImageJ and Prism 8 (GraphPad) software. The data were fitted with the Hill equation with a Hill coefficient of 2-2.5. All the nucleic acids used in this study are listed in Supplementary Table S1.

### Co-purification nucleic acids

Isolation of co-purification nucleic acids was carried out as described in ref. (18) with minor modifications. Briefly, CaCl2 and proteinase K (Zomanbio) were added to final concentrations of 5 mM and 0.5 mg/ml to 2 mg of purified *Mbp*Ago in Buffer B. The sample was incubated for 50 min at 55 °C. The nucleic acids were separated from the organic fraction by adding Roti-phenol/chloroform/isoamyl alcohol pH 7.5–8.0 in a 1:1 ratio. The top layer was isolated and nucleic acids were precipitated using ethanol precipitation by adding 99% ethanol in a 1:2 ratio supplied with 0.5% linear polymerized acrylamide as a carrier. This mixture was incubated for 16 h at −20°C and centrifuged in a table centrifuge at 13,000 rpm for 30 min. Next, the nucleic acid pellet was washed with 500 μl of 70% ethanol and solved in 50 μl nuclease-free water. The purified nucleic acids were treated with either 100 μg/ml RNase A (Thermo Fisher Scientific), 2 units DNase I (NEB), or both for 1 h at 37°C and resolved on a denaturing urea polyacrylamide gel (20%) and stained with SYBR Gold.

## RESULTS

### *Mbp*Ago prefers to cleave RNA rather than DNA with small DNA guides at ambient temperature

*Mbp*Ago (accession number WP_008504757.1 in the NCBI protein database) is distantly related to other characterized eAgos and pAgos (sequence identity < 20%) (Figure 1A and S1D) and contains the canonical catalytic tetrad in the PIWI domain (residues D566, E601, D635, and D768) (Supplementary Figure S1D). To study the biochemical properties and *in vivo* functions of *Mbp*Ago, we expressed and purified *Mbp*Ago. Codon-optimized gene encoding *Mbp*Ago was chemically synthesized and cloned into pET28a plasmid (Supplementary Figure S1A). In addition to the wild-type protein, we obtained its catalytically inactive variant (*Mbp*Ago_DM) with substitutions of two out of four catalytic tetrad residues (D566A/D635A) (Supplementary Figure S1D). The protein was expressed in *E. coli* and purified using Ni-NTA-affinity and size-exclusion (see Supplementary Figure S1 and Materials and Methods for details). Examination of purified *Mbp*Ago showed high purity of the samples (Supplementary Figure S1).

**Figure 1.**
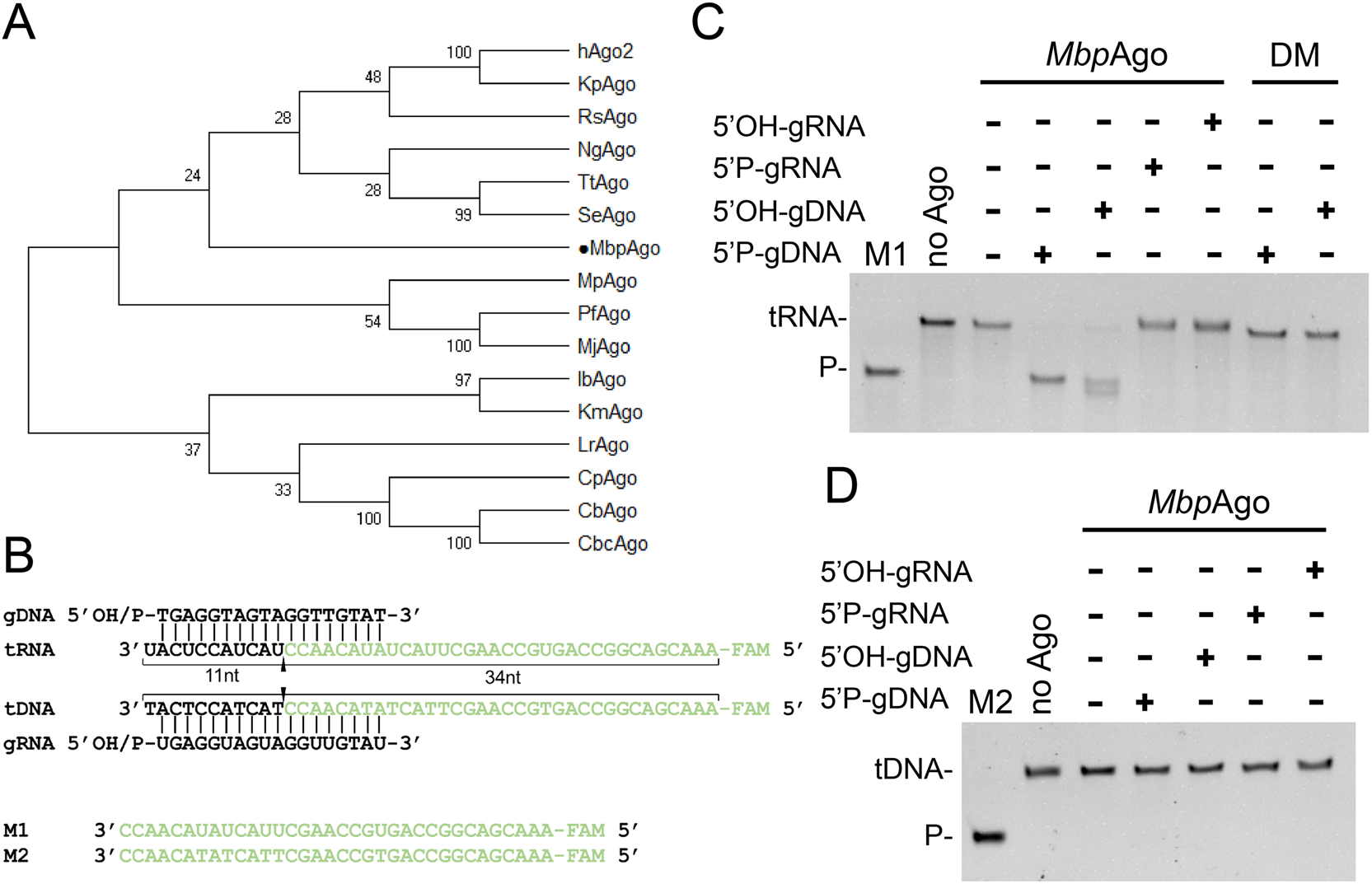
*Mbp*Ago exhibits DNA-guided RNA endonuclease activity at 37°C. (A) Maximum likelihood phylogenetic tree of characterized Ago proteins. (B) Guide and target oligonucleotides. DNA guides and RNA targets were used in most experiments. Black triangle indicates the cleavage site. (C) *Mbp*Ago exhibits DNA-guided RNA endonuclease activity. (D) *Mbp*Ago exhibits no DNA cleavage activity. Positions of the cleavage products (P) are indicated on the left of the gels. *Mbp*Ago, guides and targets were mixed in a 4:2:1 molar ratio (800 nM *Mbp*Ago preloaded with 400 nM guide, plus 200 nM target) and incubated for 30 min at 37°C. Catalytic mutant *Mbp*Ago_DM (DM) was used as a control. Lanes M1 and M2 contain chemically synthesized 34-nt RNA and DNA corresponding to the cleavage products of target RNA and target DNA, respectively.

We first studied the nucleic acid specificity of *Mbp*Ago in an *in vitro* cleavage assay using synthetic fluorescently labeled oligonucleotides targets (Figure 1B). *Mbp*Ago was loaded with 18 nt DNA or RNA guides containing a 5’phosphate or 5’hydroxyl group at 37°C for 10 min followed by the addition of complementary 5’end FAM-labeled 45-nt long single-stranded DNA or RNA targets (Figure 1B, Supplementary Table S1). After incubation for 30 min at 37°C, the cleavage products were resolved by 20% denaturing gel (Figure 1C and 1D). *Mbp*Ago uses both 5’phosphorylated DNA guide (5’P-gDNA) and 5’hydroxylated DNA guide (5’OH-gDNA) to cleave RNA target (tRNA) (Figure 1C). Unexpectedly, no DNA target (tDNA) cleavage was observed (Figure 1D), and only very weak tDNA cleavage was observed even after incubation for 6 h (Supplementary Figure S1E). With the RNA guide, we did not observe any *Mbp*Ago-mediated cleavage, neither of DNA nor RNA targets, even after incubation for 12 h (Figure 1C, 1D and Supplementary Figure S1E). This DNA-guided RNA target preference has not been observed in other eAgos or pAgos homologs, for eAgos prefer to use RNA guides to target RNA and pAgos prefer to use DNA guides to target DNA. Cleavage of the target strand by *Mbp*Ago occurs after the 10th nucleotide counting from the 5’end of the 5’P-gDNA, consistent with previously characterized Ago homologs (Supplementary Figure S1F and S1G) (14). When guided with 5’OH-gDNA, target cleavage not only occurred between target position 10’-11’ relative to the guide 5’end but also occurred at 1-2 nucleotides downstream of the canonical cleavage site (Supplementary Figure S1F and S1G), which is similar to that catalyzed by *Cb*Ago (17) and *Km*Ago (20). No cleavage activity was detected for a catalytically dead *Mbp*Ago variant with substitutions of two of the tetrad residues (D566A/D635A) (Figure 1C). In summary, in contrast to previously characterized pAgos that prefer to target DNA, *Mbp*Ago prefers to target RNA rather than DNA and can potentially be exploited for RNA manipulation.

### *Mbp*Ago uses 5’OH-gDNA and 5’P-gDNA to cleave RNA with almost the same efficiency and is active over a wide range of temperatures

The majority of pAgos strongly prefer 5’-phosphorylated nucleic acid guides (5,17,20). As shown in Figure 1C and Supplementary Figure S2A, *Mbp*Ago can use 5’P-gDNA and 5’OH-gDNA to cleave almost all RNA target in 30 min, but with different precision. We next tested RNA target cleavage kinetics and the results showed that *Mbp*Ago uses 5’OH-gDNA and 5’P-gDNA to cleave RNA with almost the same efficiency (Figure 2A). To explore the temperatures range at which *Mbp*Ago is active, we tested the temperature-dependent of RNA cleavage activity. *Mbp*Ago bound with 5’P-gDNA had comparable levels of RNA cleavage activity between 25 and 65°C, and retained good activity to cleave RNA at 4-20°C. For 5’OH-gDNA, the activity was enhanced with increasing temperature from 4 to 50°C, and then decreased rapidly with higher temperature (Figure 2B, Supplementary Figure S2B and S2C), suggesting that interactions with the 5’-phosphate may be essential to stabilize the binary *Mbp*Ago-guide complex at elevated temperatures (17).

**Figure 2.**
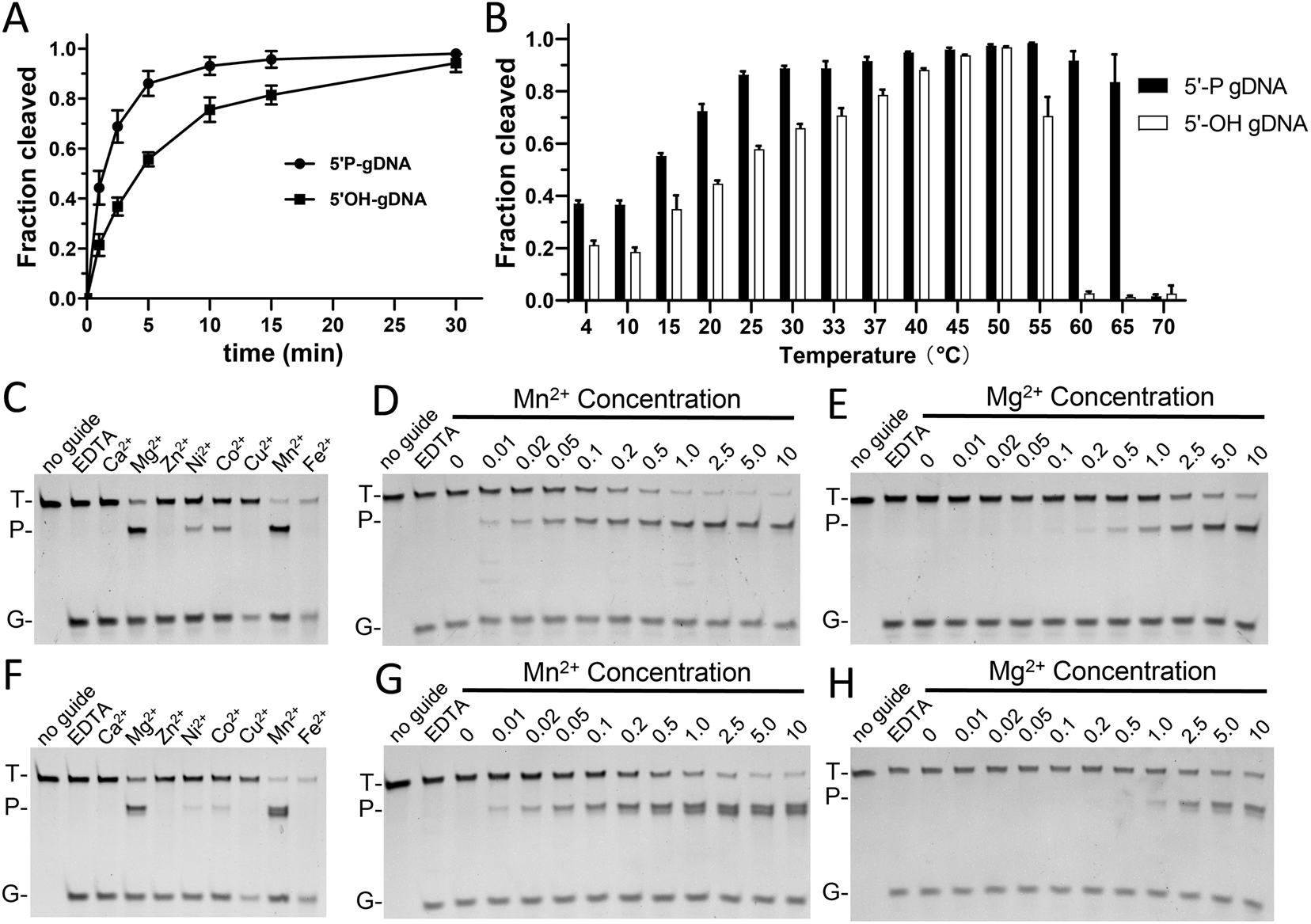
*Mbp*Ago can utilize both 5’-phosphorylated and 5’-hydroxylated DNA guides to cleave RNA targets at 37°C. (A) Kinetics of nucleic acid cleavage by *Mbp*Ago measured with FAM-labeled RNA targets. (B) Temperature dependence of RNA cleavage by *Mbp*Ago. The assay was performed for 15 min at indicated temperatures. (C) 5’-phosphorylated DNA-guided RNA cleavage by *Mbp*Ago with various divalent cation. (D) and (E) Effects of Mn^2+^ concentration or Mg^2+^ concentration on RNA cleavage activity mediated by 5’-phosphorylated DNA guides. (F) 5’-hydroxylated DNA-guided RNA cleavage by *Mbp*Ago with various divalent cation. (G) and (H) Effects of Mn^2+^ concentration or Mg^2+^ concentration on RNA cleavage activity mediated by 5’-hydroxylated DNA guides. Positions of the guides (G), targets (T) and cleavage products (P) are indicated on the left of the gels. Experiments in (C) (D) (E) (F) (G) (H) were carried out for 30 min at 37°C.

### Divalent cations and guide lengths affect the cleavage of *Mbp*Ago

We next tested which divalent cations *Mbp*Ago can utilize to mediate DNA-guided RNA target cleavage. *Mbp*Ago can utilize Mn^2+^, Mg^2+^, Ni^2+^ and Co^2+^ as cation, with Mn^2+^ being a better cation than other cations (Figure 2C and 2F). Titration of Mn^2+^ ions showed that *Mbp*Ago was active between 0.01 and 10 mM Mn^2+^ (Figure 2D and 2G), regardless of whether the guide is 5’P-gDNA or 5’OH-gDNA. Titration of Mg^2+^ ions showed that *Mbp*Ago was active between 0.1 and 10 mM Mg^2+^ (Figure 2E and 2H), regardless of whether the guide is 5’P-gDNA or 5’OH-gDNA.

We further investigated *Mbp*Ago cleavage efficiency using DNA guides of different lengths, revealing that *Mbp*Ago was most active with 15-20 nt DNA guides, with a lower efficiency observed with longer or shorter guides (Figure 3A). Similar to that of *Km*Ago, the cleavage position was shifted if shorter (11-13 nt) and longer (21-40) 5’P-gDNAs were used (20,21). Interestingly, the cleavage only occurred between the 10th and 11th guide positions when using 15-17 nt long 5’OH-gDNA, while the cleavage position was shifted if shorter (10-14 nt) or longer (18-30 nt) 5’OH-gDNA were used (Figure 3A). In conclusion, the length of the guide has no great influence on the RNA cleavage activity but has an influence on the cleavage precision.

**Figure 3.**
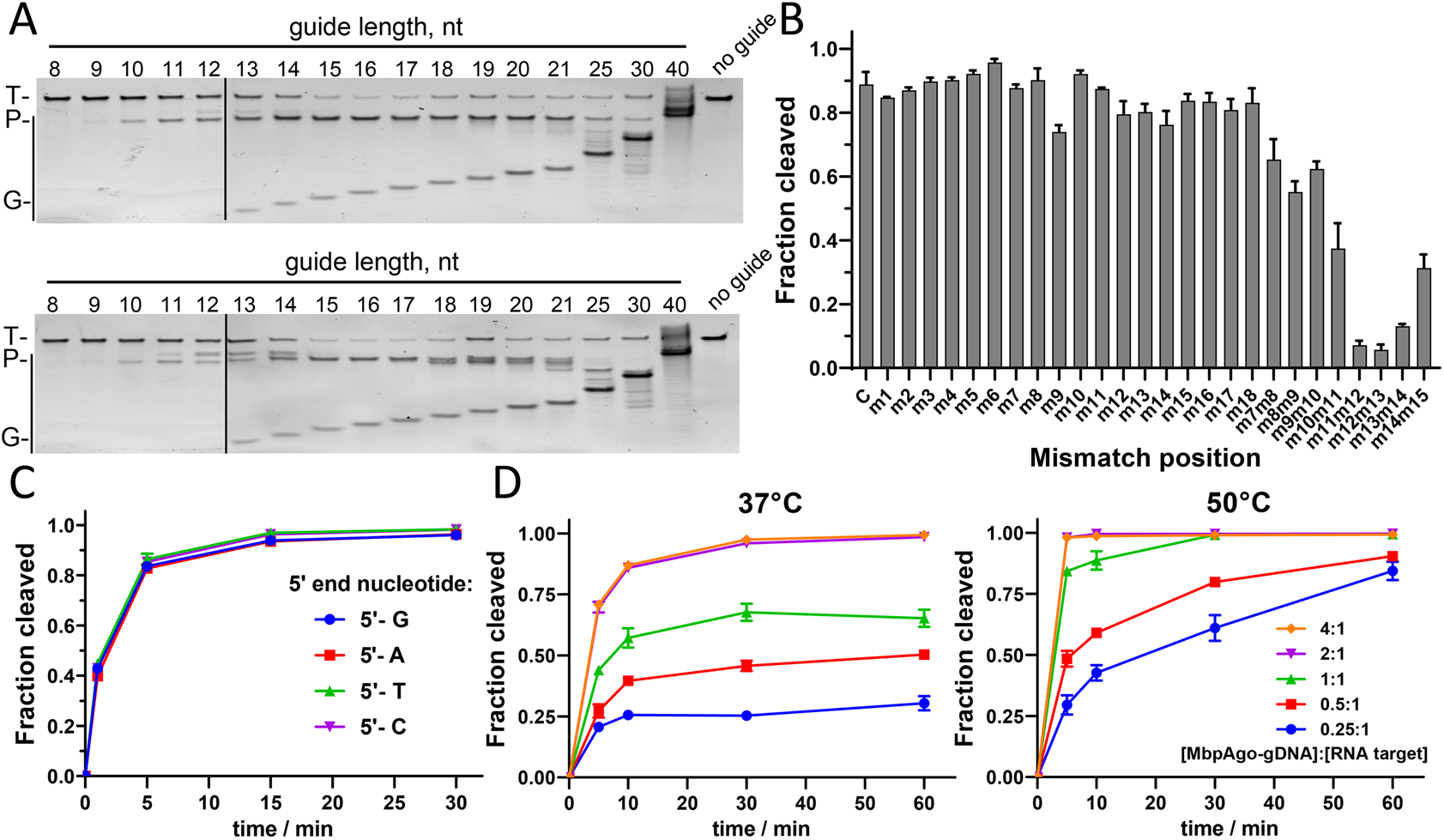
Effects of the guide on RNA cleavage by *Mbp*Ago. (A) Cleavage assay with 5’-phosphorylated (Upper panel) and 5’-hydroxylated DNA guides (Lower panel) of varied length. (B) Effects of guide-target mismatches on RNA cleavage by *Mbp*Ago. (C) Preferences for the 5’-guide nucleotides. (D) Quantified data of the *Mbp*Ago-mediated DNA-guided RNA cleavage turnover experiments using 2 pmol RNA target and increasing concentrations of *Mbp*Ago-gDNA (0.5-8 pmol) at 37°C (left panel) and 50°C (right panel). Experiments in (B) (C) (D) were carried out with 5’-phosphorylated DNA guide.

To test whether *Mbp*Ago is a single-turnover enzyme or a multiple-turnover enzyme, we monitored the cleavage assays in a time course using variable [*Mbp*Ago-gDNA]: RNA target ratios (Figure 3D) at 37°C and 50°C, respectively. We observed a rapid burst of activity during the first minute, but this stage was followed by a slow steady-state and finally only a fraction of the target was cleaved at 37°C during the 60 min course of the reaction (Figure 3D, left panel; Supplementary Figure S2F). In contrast, when the reaction was performed at 50°C almost all RNA was cleaved during 60 min, indicative of multiple-turnover reaction (Figure 3D, right panel; Supplementary Figure S2G). In conclusion, while *Mbp*Ago function as a multiple-turnover enzyme, its steady-state rate may be limited by product release and exchange of target molecules like that of *Cb*Ago and *Km*Ago (18,21).

### Effects of 5’ guide nucleotide and guide-target mismatches on target cleavage

Previous studies demonstrated that the first nucleotide in the guide strand is bound in the MID pocket of Ago, and many studied eAgos and pAgos have a certain preference for it (5). To determine if *Mbp*Ago has specificity for the first nucleotide of the guide, we also tested four guide variants with different first nucleotides but otherwise identical sequences. Similar to that of *Cb*Ago and *Km*Ago, we observed no changes in the cleavage efficiency and rate when *Mbp*Ago was loaded with 5’phosphorylated DNA guides with different first nucleotides (Figure 3C and Supplementary Figure S2E) (17,21).

Mismatches between the guide and target have effects on the cleavage activities of Ago proteins (17,20,29). To determine the sequence specificity of *Mbp*Ago, we analyzed the effects of mismatches between the guide and target strands on its RNA cleavage activity. We designed a set of DNA guides, each containing a single-nucleotide or dinucleotide mismatches at a certain position (Supplementary Table S1), and tested them in the RNA cleavage reaction with *Mbp*Ago (Figure 3B and Supplementary Figure S2D). Despite mismatch at position 9 decreased the cleavage efficiency, other single-nucleotide mismatches had no significant effect on cleavage. However, dinucleotide mismatches at position 10-15 dramatically reduced target cleavage, which suggested that *Mbp*Ago can potentially be programmed for specific cleavage of desired target sequence.

### Binding analysis of guides and targets by *Mbp*Ago

Our results indicated that *Mbp*Ago could use 5’OH DNA guides to cleave the target RNA with similar efficiency in comparison to 5’P DNA guides in figure 2A. Previously structural and bioinformatic studies showed that the MID and PAZ domain anchors the 5’end of a guide by a conserved six-amino-acid motif, and the first two residues of this motif might be important for the function of different groups of pAgos (8). Multiple sequence alignment of the 5’end guide binding pocket of the MID domain from *Mbp*Ago with several other characterized Ago proteins showed that the first two residues of most characterized Ago proteins are YK, while the first two residues of *Mbp*Ago are HK (Supplementary Figure S3A). With the structure of *Cb*Ago with its 5’P DNA guide and DNA target (PDB ID: 6qzk, identity 18.8%) as a template, we used the SWISS-MODEL portal to build the structure of *Mbp*Ago, which displays four typical domains of pAgo protein structure (Figure 4A). The 3D model aliged to *Cb*Ago revealed that first two amino acid residues in the specific motif involved in interaction with the guide 5’-end, which consistent with the multi-sequence alignment results (Figure 4A and Supplementary Figure S3A). The imidazole of the H482 could stack on the first base at the guide 5’-end in the MID domain of *Mbp*Ago like the Y472 in the *Cb*Ago. For 5’OH DNA guide, the presence of potential contacts between the MID Domain and the guide 5’-end might compensate for the loss of interactions with the 5’-phosphate in *Mbp*Ago (17).

**Figure 4.**
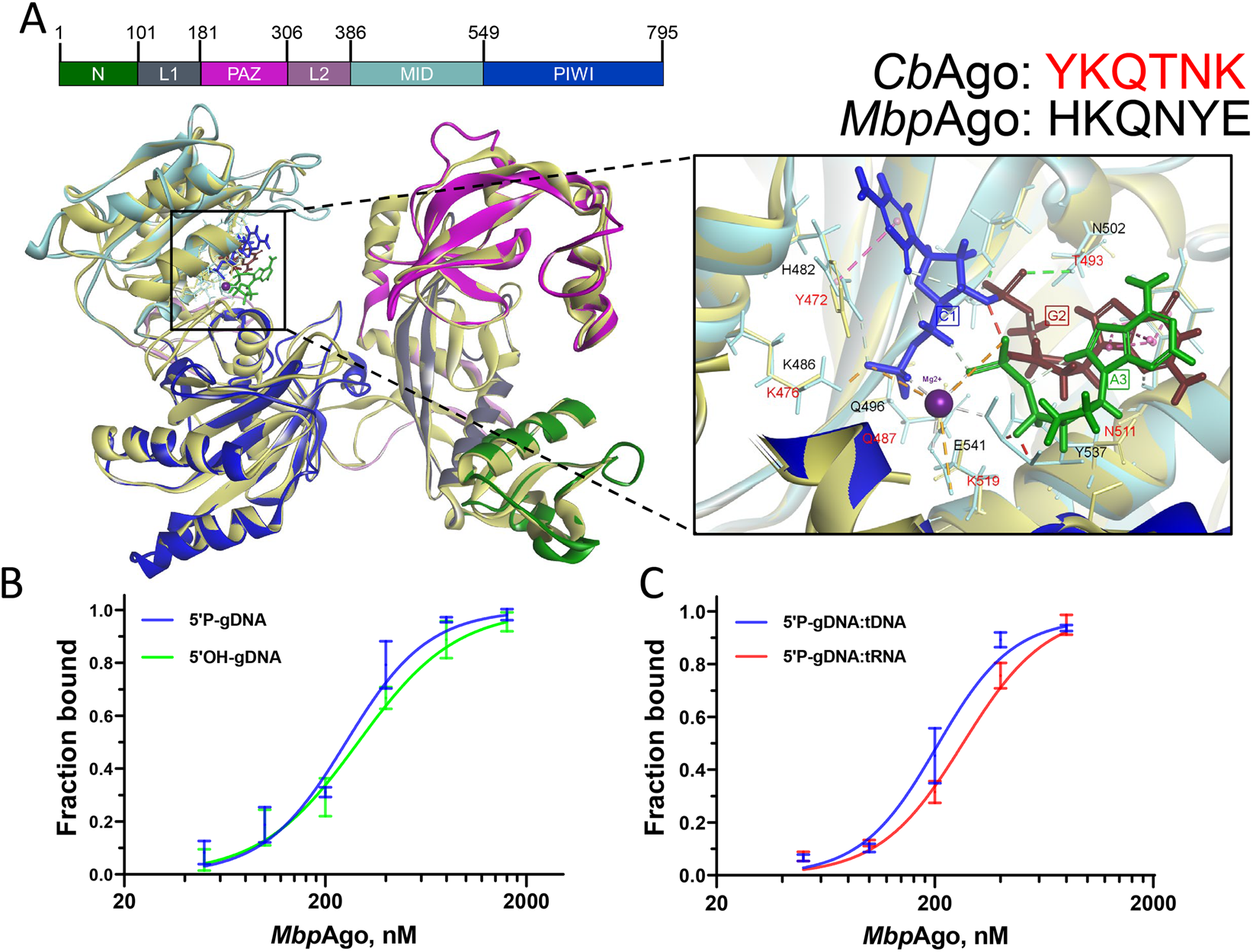
Binding analysis of guides and targets by *Mbp*Ago. (A) (left panel) A 3D model of the *Mbp*Ago aligned to the structure of *Cb*Ago in complex with gDNA and tDNA (with bound Mg^2+^ ion; PDB: 6QZK). Domains of *Mbp*Ago are coloured according to the coloured domain architecture of *Mbp*Ago with residues numbered and *Cb*Ago is coloured with light yellow. The model was built using the SWISS-MODEL portal. (right panel) Amino acid residues of the conserved MID-domain motif (shown for *Mbp*Ago and *Cb*Ago above the structure) and Mg^2+^ ion (purple) involved in interactions with the first nucleotide (blue) and the second nucleotide (deep red) of guide are highlighted. Elements of the secondary structure and amino residues specific to *Mbp*Ago and *Cb*Ago are shown in cyan and light yellow, respectively. (B) Binding of 18 nt guide by *Mbp*Ago. The fraction of bound guide was plotted against protein concentration and fitted using the model of specific binding with the Hill slope. *Mp*Ago binds the 5’P-gDNA and 5’OH-gDNA with average Kd values of 253.7 ± 23.3 nM and 291.8 ± 25.2 nM, respectively. Data are represented as the mean ± SD from three independent experiments. (C) Binding of guide: target duplex by *Mbp*Ago. The fraction of bound guide: target duplex was plotted against protein concentration and fitted using the model of specific binding with the Hill slope. *Mbp*Ago binds the 5’P-gDNA: tRNA duplex and 5’OH-gDNA: tDNA duplex with average Kd values of 203.9 ± 27.2 nM and 269.2 ± 36.5 nM, respectively. Data are represented as the mean ± SD from three independent experiments.

We further measured the dissociation constants (Kd) for guide binding by *Mbp*Ago using EMSA (Figure 4B and Supplementary Figure S3B). *Mbp*Ago has similar Kd values for 5’P DNA guides 253.7 nM (95% CI: 214.3 to 277.0 nM) and 5’OH DNA guides 291.8 nM (95% CI: 245.9 to 317.0 nM), which may be the reason why *Mbp*Ago is able to use both 5’P and 5’OH DNA guides for efficiently RNA target cleavage. We also measured the dissociation constants for guide: target duplex binding by *Mbp*Ago using EMSA (Figure 4C and Supplementary Figure S3C). As shown in Figure 4C and Supplementary Figure S3C, *Mbp*Ago is able to bind both 5’P-gDNA: tRNA duplex and 5’P-gDNA: tDNA duplex with comparable affinity, which suggested that the weak DNA target cleavage activity may be not due to target binding affinity. Furthermore, we analyzed the nucleic acids that co-purify with *Mbp*Ago after expression in *E. coli* (Supplementary Figure S1H). Small DNA was not observed in RNase A-treated samples, while RNAs of undefined length were observed in DNase I-treated samples. They are suspected to be non-specifically bound nucleic acids, as has previously been described for purification of *Tt*Ago and *Pf*Ago (3,4).

### *Mbp*Ago can use 5’P-gDNA and 5’OH-gDNA to cleave highly structured SARS-CoV-2 RdRp RNA

The coronavirus disease 2019 (COVID-19) pandemic, caused by the severe acute respiratory syndrome coronavirus 2 (SARS-CoV-2) virus, has highlighted the need for antiviral approaches. The SARS-CoV-2 virus belongs to a family of positive-sense RNA viruses, and Abbott et al reported a CRISPR-Cas13-based strategy for viral inhibition that can effectively degrade RNA from SARS-CoV-2 sequences (30). To explore whether *Mbp*Ago can cleave RNA efficiently, we designed a set of gDNAs corresponding to different regions of the highly conserved RNA-dependent RNA polymerase (RdRP) gene, which maintains the proliferation of SARS-CoV-2. We hypothesized that secondary structure of the SARS-CoV-2 RdRp RNA, as predicted by the NUPACK algorithm (Figure 5A) (31). *Mbp*Ago was loaded with gDNAs corresponding to different regions of SARS-CoV-2 RdRp RNA that either maintain single-stranded conformation or are involved in extensive base-pairing (Figure 5A and 5B). Cleavage products were detected with all 5’P-gDNAs and 5’OH-gDNAs when reactions were performed with 5 mM Mn^2+^ (Figure 5C and 5D). Moreover, *Mbp*Ago can cleave SARS-CoV-2 RdRp RNA at the expected position for all sites with 5 mM Mg^2+^, albeit to a little less efficient (Supplementary Figure S4A and S4B).

**Figure 5.**
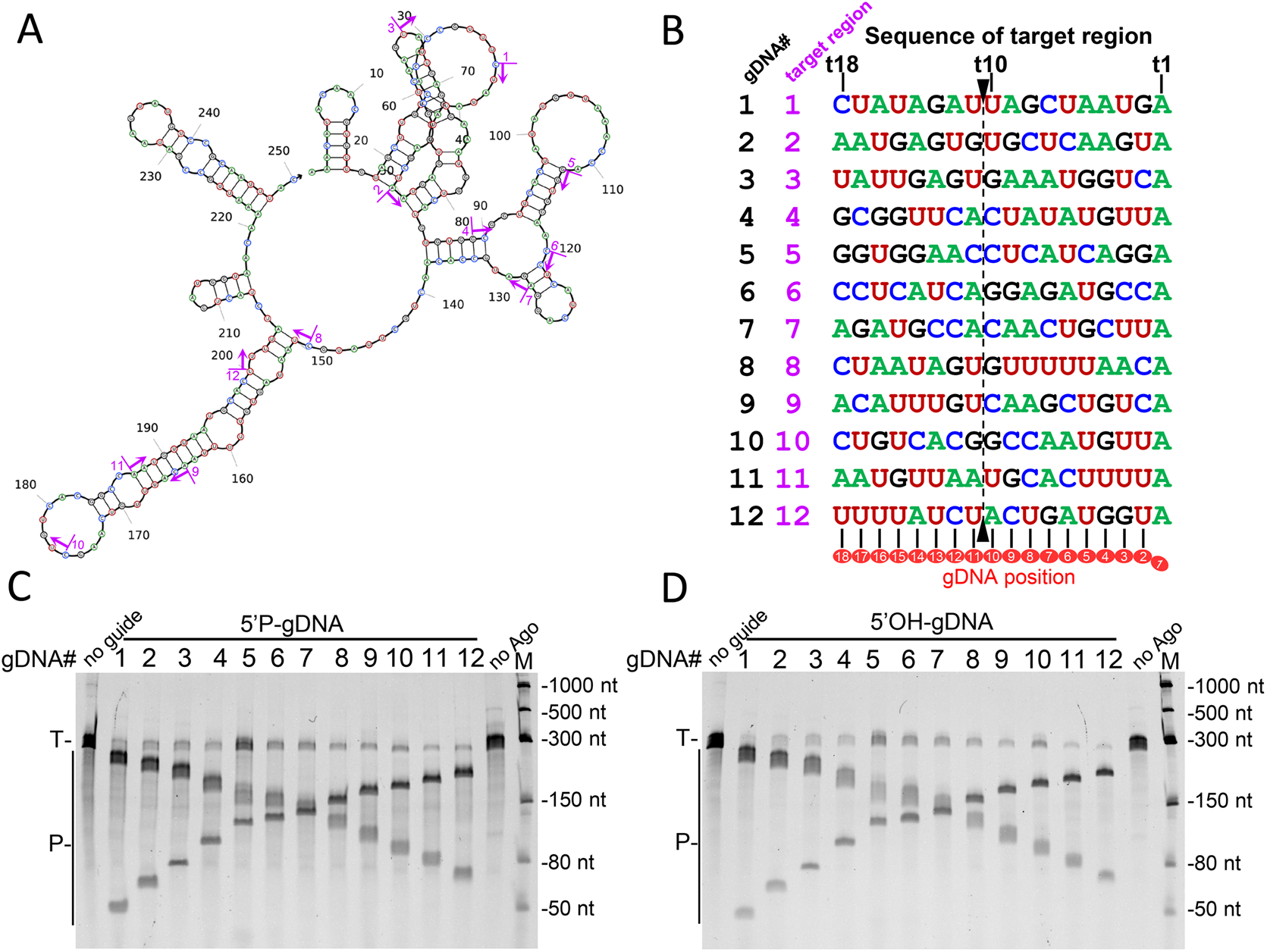
Cleavage of highly structured SARS-CoV-2 RdRp partial RNA by *Mbp*Ago-gDNA complex. (A) Secondary structure of the SARS-CoV-2 RdRp partial RNA used in the experiments predicted by NUPACK. DNA guides targeting the different target regions of the SARS-CoV-2 RdRp RNA are indicated by purple bars and arrows (1–12). (B) Schematic overview of the designed target regions in the SARS-CoV-2 RdRp RNA. Dotted line flanked by black arrowheads indicates cleavage site. (C) Analysis of the cleavage products obtained after incubation of 5’P-gDNA-*Mbp*Ago complex with SARS-CoV-2 RdRp RNA. (D) Analysis of the cleavage products obtained after incubation of 5’OH-gDNA-*Mbp*Ago complex with SARS-CoV-2 RdRp RNA. Experiments in (C) and (D) were carried out with 5 mM Mn^2+^.

## DISCUSSION

Most characterized mesophilic pAgos proteins were shown to strongly prefer DNA targets (17–20), and *Ng*Ago (*Natronobacterium gregory*) was reported to only degrade RNA targets by an unknown mechanism with poor accuracy (32). In this study, we have characterized a novel prokaryotic Argonaute protein from the psychrotolerant bacteria *Mucilaginibacter paludis* (33). Though *Mbp*Ago prefers to use the DNA guides like most characterized pAgos, *Mbp*Ago has an unusually preference for cleaving RNA targets with high precision and efficiency at moderate temperatures but has very weak activity in cleaving DNA targets. We have shown that *Mbp*Ago can target RNA under a wide range of reaction conditions. The efficiency and accuracy of cleavage are modulated by temperature, divalent ions, the phosphorylation and length of the DNA guides and its complementarity to the RNA targets.

In contrast to other studied Agos, *Mbp*Ago is able to use 5’OH guides for RNA targets cleavage with almost the same efficiency as 5’P guides. Except for several pAgos, including *Mp*Ago, exclusively bind 5’OH guides (13), the majority of eAgos and pAgos were shown to use 5’P guides for RNA cleavage, and multiple interacts between the 5’P group and the MID domain are observed in the structure of some Ago-guide complexes (5,18). A bioinformatic study revealed several subtypes of the MID domain with substitutions of key residues involved in interactions with the 5’end of guide molecule (8). The MID domain of most Agos, including *Cb*Ago, *Rs*Ago, and *Km*Ago, contains YK subtype, while *Mbp*Ago contains HK subtype (Supplementary Figure S3A). As far as we know, *Mbp*Ago might be the first studied pAgo with HK subtype in the motif. Furthermore, homology-based structural modeling suggests that interactions of the guide 5’end with the MID pocket are overall very similar for *Cb*Ago and *Mbp*Ago (Figure 4A). Therefore, interactions with other parts of the guide may be important for stabilize the 5’OH guide in *Mbp*Ago. Moreover, *Mbp*Ago binds 5’P (~253.7 nM) and 5’OH (~291.8 nM) DNA guides with comparable binding affinities. The affinities are much lower in comparison with *Cb*Ago (~6.2 pM), *Mj*Ago (~3 nM), and *Rs*Ago (~42.6 nM) but similar to hAgo2 (~565 nM) (17,34–36). Furthermore, both DNA targets and RNA targets can be bound to *Mbp*Ago with similar binding affinities. The conformational changes of *Mbp*Ago in the process of binding to the DNA target may not make the nucleic acid duplex and the catalytic cleft close enough, resulting in the weak DNA cleavage activity (29,37). Thus, structural studies will be necessary to determine the structural basis of the apparent preference for RNA targets. We were unable to co-purify DNA guides from *Mbp*Ago heterologously expressed in *E. coli*. Possibly *Mbp*Ago is not able to generate enough guides for its poor DNA cleavage activity or requires host factors for guide generation and/or loading (4). Future studies should focus on guides associated with *Mbp*Ago expressed in *Mucilaginibacter paludis*.

*Mbp*Ago is able to cleave RNA targets at a wide range of temperatures and still has good RNA cleavage activity at 4-20°C, which may be due to *Mbp*Ago is from a psychrotolerant bacteria. This suggests pAgos with good activity at mesophilic temperatures could be also found from psychrotolerant and psychrophilic bacterias. *Mbp*Ago preferentially uses 15-18 nt long DNA guides and is mostly active as RNA endonuclease in the presence of Mg^2+^ or Mn^2+^, similarly to previously studied pAgos. Interestingly, we have shown that the efficiency and the position of cleavage can be modulated by the length of the guide. While the cleavage site is shifted if shorter or longer 5’OH guides are used, the cleavage position is not shifted if 15-17 nt long 5’OH guides are used. That means we can use *Mbp*Ago to cleave RNA targets with high efficiency and precision by choosing 5’OH-gDNA of appropriate length. Moreover, the cleavage site is also shifted if 18 nt 5’OH guides are used in some pAgos, including *Cb*Ago, *Cp*Ago, *Ib*Ago, *Lr*Ago and *Km*Ago. Choosing 5’OH guides of appropriate length could use these pAgos to cleave targets with high efficiency and precision. *Mbp*Ago has no obvious preference for the 5’end nucleotide of a guide, which is similar to several other pAgos, including *Km*Ago and *Cb*Ago (17,20). Importantly, the cleavage activity of *Mbp*Ago is greatly affected when introducing mismatches with the RNA target. We have shown that the cleavage efficiency can be regulated by mismatches in the central and 3’supplementary regions of the guide.

Finally, we have demonstrated that *Mbp*Ago can efficiently cleave highly-structured RNA targets using both 5’P-gDNAs and 5’OH-gDNAs in the presence of Mg^2+^ or Mn^2+^. Previously, eAgo from the budding yeast *Kluyveromyces polysporus* and pAgo from the mesophilic bacteria *Kurthia masiliensis* were used for highly structured RNA cleavage (21,38). Different from their findings, the cleavage efficiency of *Mbp*Ago does not greatly depend on the secondary structure at 37°C. Therefore, *Mbp*Ago can be used for cleavage of complex RNA targets independently of the secondary structure formation. Furthermore, using *Mbp*Ago to cleave RNA is more convenient and cost-effective, for the synthesis of 5’OH DNA guides is easier and more inexpensive than 5’P DNA guides. Thus, *Mbp*Ago can potentially be applied in RNA-centric *in vivo* and *in vitro* methods such as nucleic acid detection, RNA targeting, and antiviral (30,39). In conclusion, we have demonstrated that *Mbp*Ago is a unique programmable nuclease that has a strong preference for RNA targets and can potentially be widely used in the field of nucleic acid biotechnology.

