## Supplemental Table and figure for "A programmable pAgo nuclease with RNA target preference from the psychrotolerant bacteria *Mucilaginibacter paludis*"

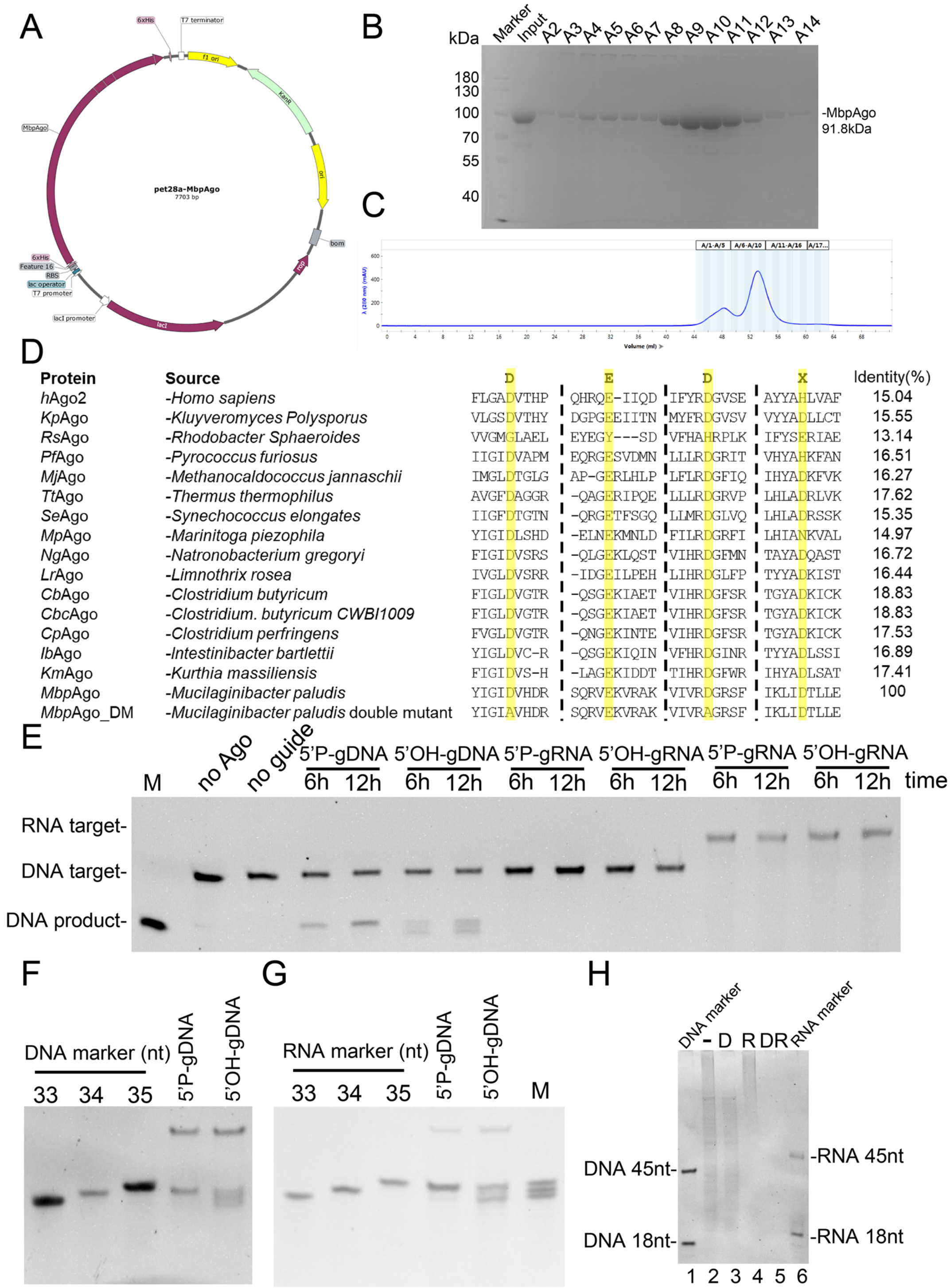


Supplementary Figure S1. *Mbp*Ago harbors a catalytic DEDD tetrad. (A) Schematic representation of the pET28a based expression vector of *Mbp*Ago. (B) The purity of the purified *Mpb*Ago was determined using SDS-PAGE. (C) Size exclusion diagram showing the elution peak of *Mbp*Ago. (D) Multiple sequence alignment of a part of the PIWI domain from *Mbp*Ago with several other characterized Ago proteins. (E) *Mbp*Ago shows DNA cleavage activity at 37°C if incubation is extended. (F) Determining the DNA cleavage site with non-labeled tDNA. DNA marker (33, 34, 35 nt) were partially hydrolyzed tDNA. The experiments were performed at the 4:2:1 *Mbp*Ago:guide:target molar ratio for 12 h at 37 °C. (G) Determining the RNA cleavage site with non-labeled tRNA. RNA marker (33, 34, 35 nt) were partially hydrolyzed tRNA. The experiments were performed at the 4:2:1 *Mbp*Ago:guide:target molar ratio for 30 min at 37°C. (H) Purification of *Mbp*Ago-associated nucleic acids after its expression in E. coli. Nucleic acids were treated with DNaseI (D), RNase A (R), both nucleases (DR), or left untreated (-), separated by denaturing PAGE and stained with SYBR Gold. lane1, DNA length markers; lane6, RNA length markers.

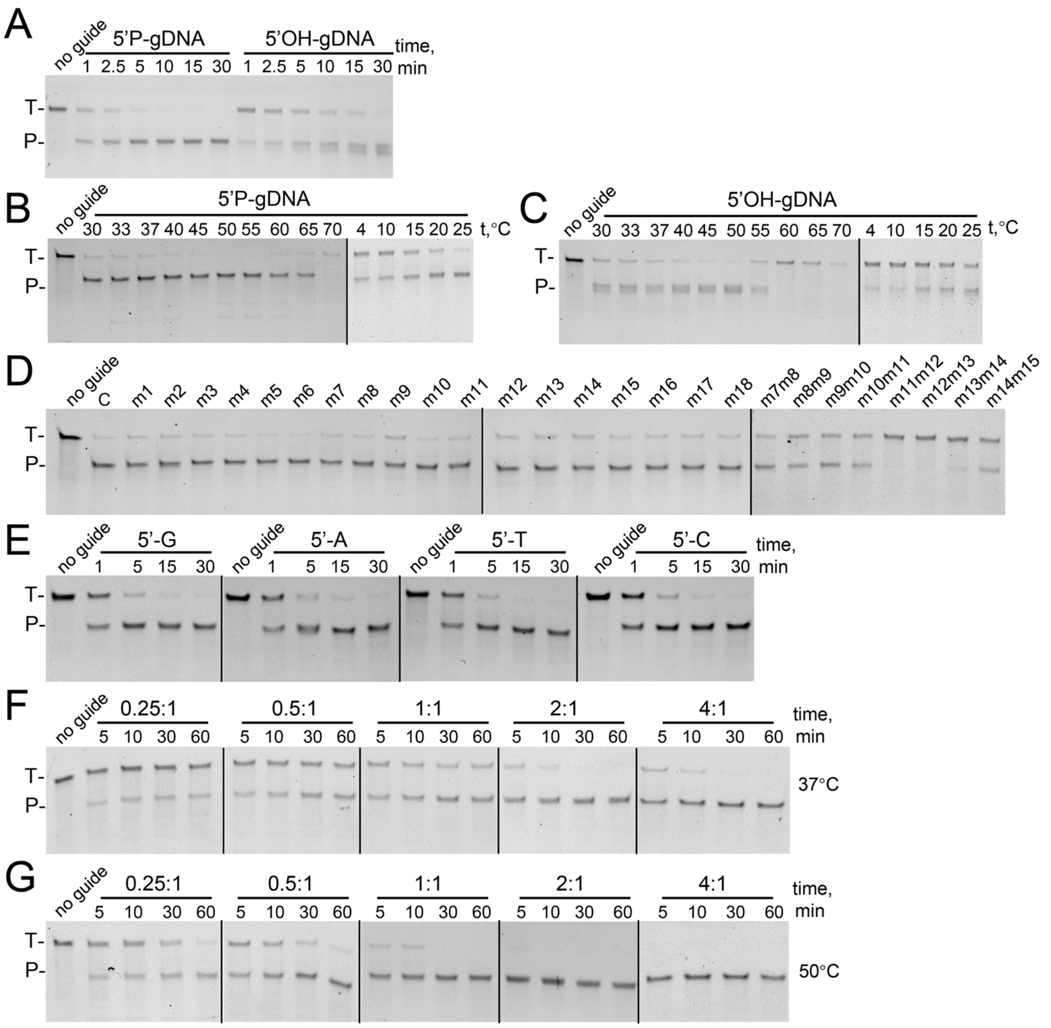


Supplementary Figure S2. Representative denaturing PAGE showing the results in Figure 2 and Figure 3. (A) Representative denaturing PAGE showing the results of RNA cleavage kinetics in the presence of 5 mM $MnCl_2$. Assays were performed in three independent replicates, and time points were taken at 1, 2.5, 5, 10, 15, 30 min. Cleavage efficiencies from three independent experiments were quantified and plotted against time (Figure 2A). (B) and (C) Representative denaturing PAGE showing effects of temperature on RNA cleavage activity using 5'P-gDNA or 5'OH-gDNA. (D) Representative denaturing PAGE showing effects of mismatches in the guide-target duplex on the slicing activity of *Mbp*Ago. (E) Representative denaturing PAGE showing the preferences for the 5'-guide nucleotides. (F) Representative denaturing PAGE showing the RNA cleavage turnover experiment at 37°C (Figure 3D). (G) Representative denaturing PAGE showing the RNA cleavage turnover experiment at 50°C (Figure 3D). All above representative denaturing PAGEs are one of three independent experiments.

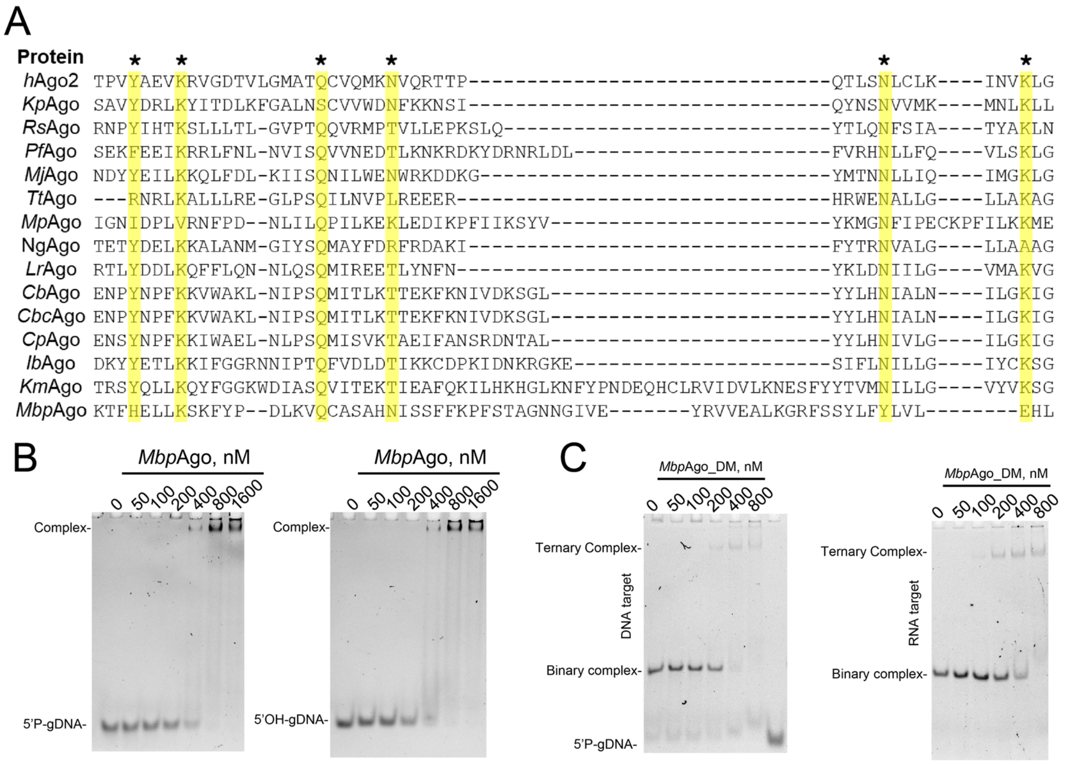


Supplementary Figure S3. Electrophoresis mobility shift assay (EMSA) of the binding of the *Mbp*Ago to guides and guide: target duplex. (A) Multiple sequence alignment of the 5'end guide binding pocket of the MID domain from *Mbp*Ago with several other characterized Ago proteins. Black asterisks are the positions of amino acid residues involved in the binding of the 5'end of a guide. (B) Representative native gel images (one of three independent experiments) of binding reactions in Figure 4B with various ratios between *Mbp*Ago and guide. (C) Representative native gel images (one of three independent experiments) of binding reactions in Figure 4C with various ratios between *Mbp*Ago and guide: target duplex.

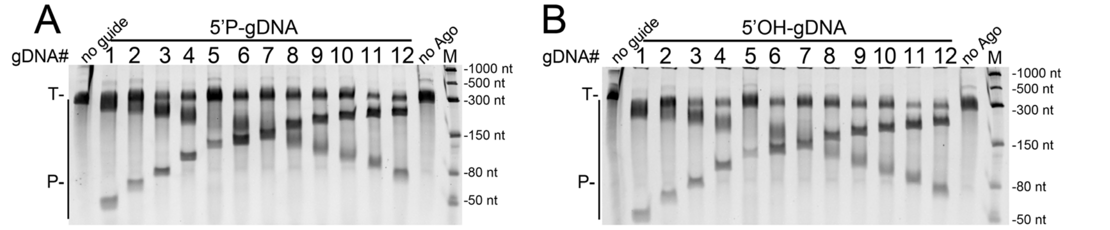


Supplementary Figure S4. Cleavage of highly structured SARS-CoV-2 RdRp RNA by *Mbp*Ago-gDNA complex with 5 mM $Mg^{2+}$. (A) Analysis of the cleavage products obtained after incubation of 5’P-gDNA-*Mbp*Ago complex with SARS-CoV-2 RdRp RNA. (B) Analysis of the cleavage products obtained after incubation of 5’OH-gDNA-*Mbp*Ago complex with SARS-CoV-2 RdRp RNA.

### Supplementary tables

**Table S1. List of let-7-derived sequences used in this study.**

| Oligonucleotide name | Sequence (5'-3') | Description |
|---|---|---|
| FAM-tDNA | FAM-AAACGACGGCCAGTGCCAAGCTTACTATACAACCTACTACCTCAT | 5’ FAM labeled T-tDNA |
| M1 | FAM-AAACGACGGCCAGTGCCAAGCTTACTATACAACC | 5’ FAM labeled 34 nt DNA |
| FAM-tRNA | FAM-AAACGACGGCCAGUGCCAAGCUUACUAUACAACCUACUACCUCAU | 5’ FAM labeled U-tRNA |
| M2 | FAM-AAACGACGGCCAGUGCCAAGCUUACUAUACAACC | 5’ FAM labeled 34 nt RNA |
| C-gDNA | CGAGGTAGTAGGTTGTAT | guide forms 5'-C pair with C-tRNA |
| T-gDNA | TGAGGTAGTAGGTTGTAT | guide forms 5'-T pair with T-tDNA/T-tRNA |
| A-gDNA | AGAGGTAGTAGGTTGTAT | guide forms 5'-A pair with A-tRNA |
| G-gDNA | GGAGGTAGTAGGTTGTAT | guide forms 5'-G pair with G-tRNA |
| 33nt DNA product | AAACGACGGCCAGTGCCAAGCTTACTATACAAC | 33 nt DNA marker |
| 34nt DNA product | AAACGACGGCCAGTGCCAAGCTTACTATACAACC | 34 nt DNA marker |
| 35nt DNA product | AAACGACGGCCAGTGCCAAGCTTACTATACAACCT | 35 nt DNA marker |
| T-tDNA | AAACGACGGCCAGTGCCAAGCTTACTATACAACCTACTACCTCAT | let-7 based 45 nt DNA target for T-gDNA/U-gRNA |
| U-gRNA | UGAGGUAGUAGGUUGUAU | guide forms 5'-U pair with T-tDNA/T-tRNA |
| U-tRNA | AAACGACGGCCAGUGCCAAGCUUACUAUACAACCUACUACCUCAU | let-7 based 45 nt RNA target for T-gDNA/U-gRNA |
| 33nt RNA product | AAACGACGGCCAGUGCCAAGCUUACUAUACAAC | 33 nt RNA marker |
| 34nt RNA product | AAACGACGGCCAGUGCCAAGCUUACUAUACAACC | 34 nt RNA marker |
| 35nt RNA product | AAACGACGGCCAGUGCCAAGCUUACUAUACAACCU | 35 nt RNA marker |
| gDNA_mm1 | AGAGGTAGTAGGTTGTAT | guide forms mismatched pair in |

| | | |
|---|---|---|
| | | position 1 with U-tRNA |
| gDNA_mm2 | TCAGGTAGTAGGTTGTAT | guide forms mismatched pair in position 2 with U-tRNA |
| gDNA_mm3 | TGTGGTAGTAGGTTGTAT | guide forms mismatched pair in position 3 with U-tRNA |
| gDNA_mm4 | TGACGTAGTAGGTTGTAT | guide forms mismatched pair in position 4 with U-tRNA |
| gDNA_mm5 | TGAGCTAGTAGGTTGTAT | guide forms mismatched pair in position 5 with U-tDNA |
| gDNA_mm6 | TGAGGAAGTAGGTTGTAT | guide forms mismatched pair in position 6 with U-tRNA |
| gDNA_mm7 | TGAGGTTGTAGGTTGTAT | guide forms mismatched pair in position 7 with U-tRNA |
| gDNA_mm8 | TGAGGTACTAGGTTGTAT | guide forms mismatched pair in position 8 with U-tRNA |
| gDNA_mm9 | TGAGGTAGAAGGTTGTAT | guide forms mismatched pair in position 9 with U-tRNA |
| gDNA_mm10 | TGAGGTAGTTGGTTGTAT | guide forms mismatched pair in position 10 with U-tRNA |
| gDNA_mm11 | TGAGGTAGTACGTTGTAT | guide forms mismatched pair in position 11 with U-tRNA |
| gDNA_mm12 | TGAGGTAGTAGCTTGTAT | guide forms mismatched pair in position 12 with U-tRNA |
| gDNA_mm13 | TGAGGTAGTAGGATGTAT | guide forms mismatched pair in position 13 with U-tRNA |
| gDNA_mm14 | TGAGGTAGTAGGTAGTAT | guide forms mismatched pair in position 14 with U-tRNA |
| gDNA_mm15 | TGAGGTAGTAGGTTCTAT | guide forms mismatched pair in position 15 with U-tRNA |
| gDNA_mm16 | TGAGGTAGTAGGTTGAAT | guide forms mismatched pair in position 16 with U-tRNA |
| gDNA_mm17 | TGAGGTAGTAGGTTGTTT | guide forms mismatched pair in position 17 with U-tRNA |
| gDNA_mm18 | TGAGGTAGTAGGTTGTAA | guide forms mismatched pair in position 18 with U-tRNA |
| gDNA_m7m8 | TGAGGTTCTAGGTTGTAT | guide forms mismatched pair in position 7 and 8 with U-tRNA |
| gDNA_m8m9 | TGAGGTACAAGGTTGTAT | guide forms mismatched pair in position 8 and 9 with U-tRNA |
| gDNA_m9m10 | TGAGGTAGATGGTTGTAT | guide forms mismatched pair in position 9 and 10 with U-tRNA |
| gDNA_m10m11 | TGAGGTAGTACGTTGTAT | guide forms mismatched pair in position 10 and 11 with U-tRNA |

| | | |
|---|---|---|
| gDNA_m11m12 | TGAGGTAGTTCCTTGTAT | guide forms mismatched pair in position 11 and 12 with U-tRNA |
| gDNA_m12m13 | TGAGGTAGTTGCATGTAT | guide forms mismatched pair in position 12 and 13 with U-tRNA |
| gDNA_m13m14 | TGAGGTAGTTGGAAGTAT | guide forms mismatched pair in position 13 and 14 with U-tRNA |
| gDNA_m14m15 | TGAGGTAGTTGGTACTAT | guide forms mismatched pair in position 13 and 14 with U-tRNA |
| 8nt T-gDNA | TGAGGTAG | 8 nt guide pair with U-tRNA |
| 9nt T-gDNA | TGAGGTAGT | 9 nt guide pair with U-tRNA |
| 10nt T-gDNA | TGAGGTAGTA | 10 nt guide pair with U-tRNA |
| 11nt T-gDNA | TGAGGTAGTAG | 11 nt guide pair with U-tRNA |
| 12nt T-gDNA | TGAGGTAGTAGG | 12 nt guide pair with U-tRNA |
| 13nt T-gDNA | TGAGGTAGTAGGT | 13 nt guide pair with U-tRNA |
| 14nt T-gDNA | TGAGGTAGTAGGTT | 14 nt guide pair with U-tRNA |
| 15nt T-gDNA | TGAGGTAGTAGGTTG | 15 nt guide pair with U-tRNA |
| 16nt T-gDNA | TGAGGTAGTAGGTTGT | 16 nt guide pair with U-tRNA |
| 17nt T-gDNA | TGAGGTAGTAGGTTGTA | 17 nt guide pair with U-tRNA |
| 19nt T-gDNA | TGAGGTAGTAGGTTGTATA | 19 nt guide pair with U-tRNA |
| 20nt T-gDNA | TGAGGTAGTAGGTTGTATAG | 20 nt guide pair with U-tRNA |
| 21nt T-gDNA | TGAGGTAGTAGGTTGTATAGT | 21 nt guide pair with U-tRNA |
| 25nt T-gDNA | TGAGGTAGTAGGTTGTATAGTAAGC | 25 nt guide pair with U-tRNA |
| 30nt T-gDNA | TGAGGTAGTAGGTTGTATAGTAAGCTTGGC | 30 nt guide pair with U-tRNA |
| 40nt T-gDNA | TGAGGTAGTAGGTTGTATAGTAAGCTTGGCACTGGCCGTC | 40 nt guide pair with U-tRNA |
| 18nt FAM-gDNA | TGAGGTAGTAGGTTGTAT-FAM | 3’ FAM labeled T-gDNA |

**Table S2. Sequences of SARS-Cov2 RdRP.**

5′-AAACATACAACGUGUUGUAGCUUGUCACACCGUUUCUAUAGAUUAGCUAAUGAGUGUGCUCAAGUAUUGAGUGAAAUGGUCAUGUGUGGCGGUUCACUAUAUGUUAAACCAGGUGGAACCUCAUCAGGAGAUGCCACAACUGCUUAUGCUAAUAGUGUUUUUAACAUUUGUCAAGCUGUCACGGCCAAUGUUAAUGCACUUUUAUCUACUGAUGGUAACAAAAUUGCCGAUAAGUAUGUCCGCAAUUUAC

**Table S3. List of gDNAs targeting SARS-Cov2 RdRP.**

| gDNA# | Sequence (5′-3′) | Target region | 5′ product length (nt) |
|---|---|---|---|
| gDNA_12 | TACCATCAGTAGATAAAA | 200-217 | 207 |
| gDNA_11 | TAAAAGTGCATTAACATT | 186-203 | 193 |
| gDNA_10 | TAACATTGGCCGTGACAG | 176-193 | 183 |
| gDNA_9 | TGACAGCTTGACAAATGT | 164-181 | 171 |
| gDNA_8 | TGTTAAAAACACTATTAG | 149-166 | 156 |

| | | | |
|---|---|---|---|
| gDNA_7 | TAAGCAGTTGTGGCATCT | 129-147 | 136 |
| gDNA_6 | TGGCATCTCCTGATGAGG | 119-136 | 126 |
| gDNA_5 | TCCTGATGAGGTTCCACC | 112-129 | 119 |
| gDNA_4 | TAACATATAGTGAACCGC | 89-106 | 96 |
| gDNA_3 | TGACCATTTCACTCAATA | 65-82 | 72 |
| gDNA_2 | TACTTGAGCACACTCATT | 51-68 | 58 |
| gDNA_1 | TCATTAGCTAATCTATAG | 36-53 | 43 |
